# Genotype fingerprints enable fast and private comparison of genetic testing results for research and direct-to-consumer applications

**DOI:** 10.1101/208025

**Authors:** Max Robinson, Gustavo Glusman

## Abstract

As genetic testing expands out of the research laboratory into medical practice as well as the direct-to-consumer market, the efficiency with which the resulting genotype data can be compared between individuals is of increasing importance.

We present a method for summarizing personal genotypes, yielding ’genotype fingerprints’ that can be derived from any single nucleotide polymorphism (SNP)-based assay and readily compared to estimate relatedness. The resulting fingerprints remain comparable as chip designs evolve to higher marker densities. We demonstrate that they support applications including distinguishing genotypes of closely related individuals by relationship type, distinguishing closely related individuals from individuals from the same background population, identification of individuals in known background populations, and de novo identification of subpopulations within a large cohort in a high-throughput manner.

An important feature of genotype fingerprints is that, while fingerprints do not preserve anonymity, they summarize individual marker data in a way that prevents phenotype prediction. Genotype fingerprints are therefore well-suited to public sharing for ancestry determination purposes, without revealing personal health risk status.

## Background

We have recently published a method for converting personal genomes into ‘genome fingerprints’ that facilitate (and greatly accelerate) their comparison (Glusman et al, 2017). Our method encodes the characteristics of pairs of consecutive single nucleotide variants (SNVs) relative to a reference, as represented in variant call format (VCF) files or structurally equivalent formats. Typically, VCF files encode only differences from the reference; genomic locations in which the individual is homozygous for the reference allele are typically not stated, achieving a more compact representation.

A different representation of genetic information enumerates all observed genotypes for a predefined set of variants of interest, typically common single-nucleotide polymorphisms (SNPs). In this format, each SNP is identified by its identifier (‘rsid’, reference SNP identifier) in the dbSNP database (Sherry et al., 2001). For each rsid, the observed genotype of the individual is stated, including those for which the individual is homozygous for the reference allele. The chromosome and coordinate of the SNP may be stated as well, relative to a version of the reference that is hopefully stated in the ‘header’ of the genotype file.

This representation is suitable for reporting the results of genotyping experiments using DNA hybridization microarrays. Due to the lower cost of DNA hybridization genotyping relative to whole-genome sequencing (WGS), exome sequencing and other forms of targeted sequencing, very large numbers of genotypes have been produced to date, using a variety of array designs [a]. Array-based genotyping is also used as a quality-control step prior to WGS. In fact, the commoditization of array-based genotyping has enabled commercial companies (including 23andMe, AncestryDNA, Family Tree DNA, and others [b]) to offer this service directly to consumers (DTC). These services typically yield results with high concordance (Imai et al., 2011) and low no-call rates (Glusman et al., 2012). Nevertheless, genotyping the same individual using different array designs can yield slightly different results, as each technology has its own biases. Even when using the same technology, reference and encoding, genotyping the same individual repeatedly can give slightly different results due to the stochastic nature of genome sequencing, batch effects, or differences in the computational pipelines used. Some companies regularly reanalyze the raw data for all customers, refining the results over time; as a result, customers who download their genotype data repeatedly over the years may have slightly differing results even from the same sample.

Many methods exist for comparing genome-wide genotypes for inferring relatedness, with varying degrees of accuracy (Ramstetter et al., 2017). Most methods are very computationally demanding and require full access to the genotype data of the individuals to be compared, potentially precluding their application to the study of samples with restricted access, or by non-specialists interested in exploring their ancestry and genealogy.

We present here a method for summarizing personal genotypes, yielding ’genotype fingerprints’ that can be readily compared to estimate relatedness. The genotype fingerprints can be computed starting from any of several chip array designs, with genome coordinates expressed relative to any reference version; the resulting fingerprints are directly comparable without further conversion. Computation on the genotype fingerprints is fast and requires little memory, enabling comparison of large sets of genomes. No individual variants or other detailed features of the personal genome can be reconstructed from the fingerprint, thereby allowing private information to be more closely guarded and protected and decoupling genome comparison from genome interpretation. Fingerprints of different sizes allow balancing the speed and accuracy of the comparisons. Due to the high value of estimating relatedness, the potential applications of genotype fingerprinting range from basic science (study design, population studies) to personalized medicine, forensics, and data privacy.

## Methods

### Overview

Our algorithm summarizes an individual’s genotype as a ‘raw’ fingerprint, which is a tally of biallelic SNPs stratified by observed alleles and by variant identifiers, and taking allele frequencies into consideration (Figure 1). We then normalize the raw fingerprint to account for systematic differences in frequency between groups. The resulting ‘normalized’ fingerprint preserves differences at the species level, e.g. between individuals from different populations. Averaging the normalized fingerprints of the individuals in a population yields a ‘population’ fingerprint, which can be subtracted from an individual’s normalized fingerprint to produce a ‘population-adjusted’ fingerprint suitable for more sensitive detection of related genomes.

**Figure 1.**
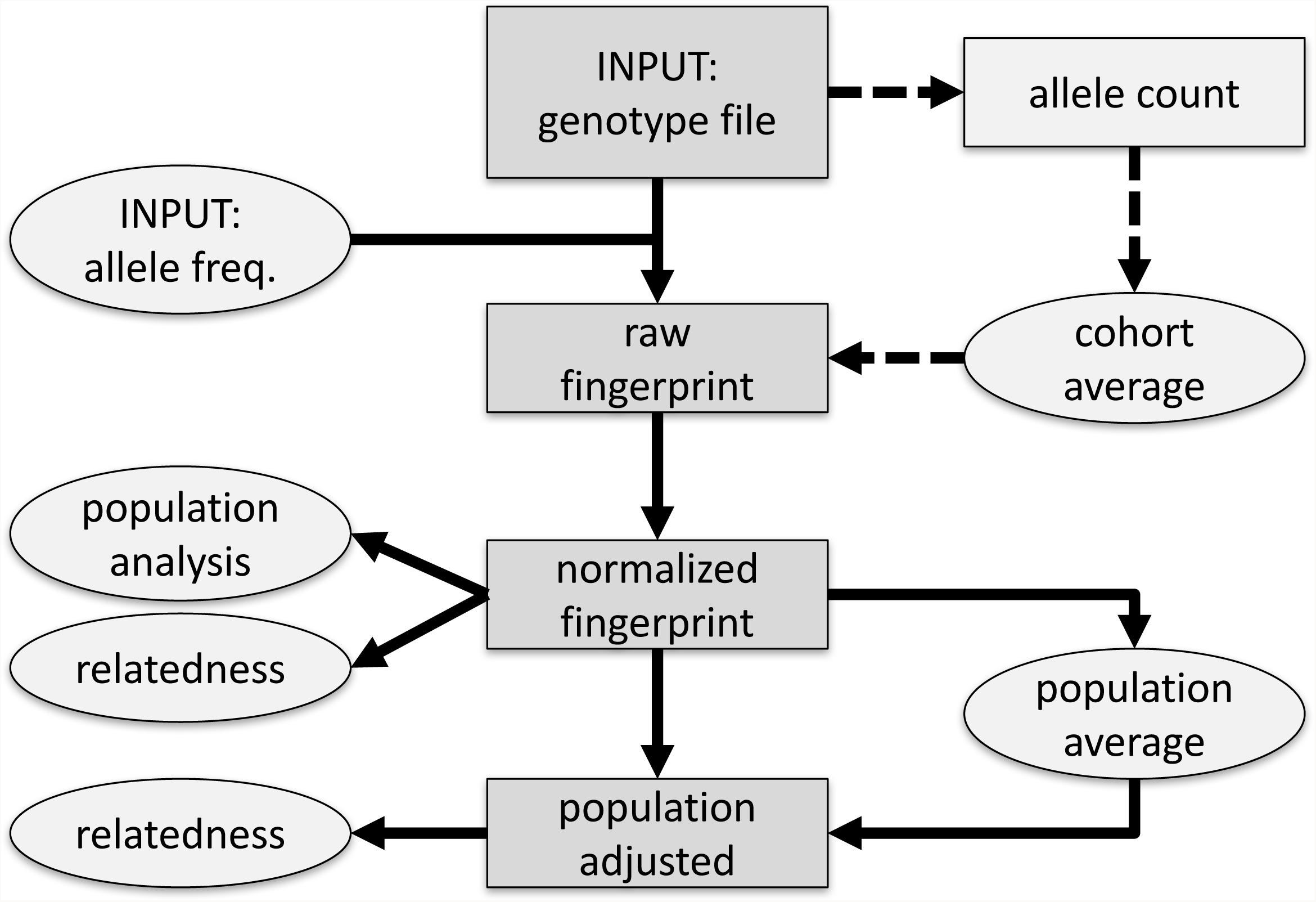
Overview of method. SNPs in the input genotype file are encoded into a table (raw) by observed alleles and rsid numerical value, taking allele frequencies into account; this can optionally be approximated by subtracting allele counts estimated from a simple model of an observed cohort (dashed arrows). The raw fingerprint is then normalized and may be adjusted to represent deviation from the center of the closest population. Rectangles and ellipses pertain to individual genotypes or to multiple genotypes, respectively; darker gray denotes the flow of information for one genotype, from the input file to the normalized and adjusted fingerprints.

### Raw fingerprints

The first stage in computing genotype fingerprints yields a ‘raw’ fingerprint, a 4 x *L* table of SNP allele counts (*L* is the main parameter of the method; defaulting to 1000). The four rows correspond to the permitted alleles A, C, G and T; variants with other possible alleles (e.g., insertions and deletions, multi-nucleotide variants) are ignored. We note for each SNP the alleles observed (both reference and alternate). We also consider the numerical component of the variant identifier (the rsid) in the dbSNP database (Sherry et al., 2001): this information determines the column in the table.

To compute a raw fingerprint:

1. Identify biallelic SNPs in autosomes as variants with alleles of length 1 in the case-insensitive alphabet [A C G T]. Ignore all other types of variants. Optionally, include only SNPs in a preselected set, e.g., 23andMe V2 and V3.
2. For each SNP, identify the numerical part of the variant identifier, e.g., 1,801,133 for rs1801133. Reduce this value modulo *L*, e.g., 133 for rs1801133 with *L*=1000. This determines the column in which the SNP will be counted.
3. For each SNP, count observations of each nucleotide on the strand represented in the reference sequence: n_A_ is the count of A alleles (0, 1, or 2), n_C_ is the count of C alleles, etc., with n_N_ = n_A_+n_C_+n_G_+n_T_ = 2.
4. In the indicated column (see 2 above), add n_A_-e_A_ to the row for A, n_C_-e_C_ to the row for C, etc., where e_X_ = n_N_ f_X_ is the expected number of X alleles and f_X_ is the expected frequency of allele X for the SNP. (Depending on the context, these frequencies may be obtained for individual SNPs, e.g. from dbSNP for human data; alternatively, they may be computed per column from all SNPs contributing to the column in an observed cohort of genotypes regardless of the nature of the cohort).

Retrieving the allele frequencies for each observed rsid requires prior knowledge and can incur in significant computational costs. It is possible to compute efficiently a data-driven approximation of the raw fingerprints as follows:

1. Compute *allele counts* as above, except in step 4 increment the value in the [4 x *L*] matrix by one for each observed allele. (In case of homozygosity, increment by two.)
2. Compute a *cohort average reference* by averaging the allele counts observed in a collection of genomes genotyped using the same array design.
3. Compute the raw fingerprint for an individual by subtracting the cohort average reference from their allele count.

### Fingerprint normalization

The second stage in computing genome fingerprints performs a normalization of the raw fingerprint to account for the different frequencies of observed alleles and numerical values of rsids. The normalization is performed in two steps:

1. Normalization by rsid value: subtract the mean and divide by the standard deviation of each column.
2. Normalization by allele: subtract the mean and divide by the standard deviation of each row.

### Adjusting fingerprints for population

We compute a population fingerprint as the average of the normalized fingerprints from the individuals in the population. These fingerprints must have been computed using the same parameter *L*. We then compute a population-adjusted fingerprint for an individual by subtracting a population fingerprint from the normalized fingerprint of that individual; both individual and population fingerprints must have been computed using the same parameter *L*.

### Fingerprint comparison

To compare two fingerprints, concatenate the rows of each fingerprint matrix into a vector and compute the Spearman correlation between the two vectors. This same procedure is appropriate for comparing two normalized fingerprints or two population adjusted fingerprints, whether adjusted to the same or different populations.

### Family analysis

We obtained 23andMe SNP chip genotype data for a family of five [c], including Mother, Father, Son, Daughter and Aunt. Son is 23andMe V2 data and the rest of the family are 23andMe V3 data. We computed normalized genotype fingerprints (*L*=5000) for the five individuals and performed all pairwise comparisons. We also extracted from these samples the lists of rsids observed in V2 and V3, for use in further analyses below.

### Population structure analysis

We studied the multi-sample VCFs in release 20130502 (filenames: ALL.chrNN.phase3_shapeit2_mvncall_integrated_v5.20130502.genotypes.vcf.gz) of the 1000 Genomes Project data set and extracted the observed genotypes, at each of the rsids in the 23andMe V2 and V3 lists, for each of the 2504 genomes in this cohort. We then computed genotype fingerprints from these deduced genotypes using *L*=500, 1000, and 5000.

We compared principal components analysis (PCA) of genome fingerprints and genotype fingerprints on this data set as follows. We identified SNPs with a minor allele frequency of 5% or more, removed SNPs in complete linkage disequilibrium with a SNP to the left (i.e. a smaller chromosomal position), retained 5% at random (298,454 SNPs) and counted occurrences of the minor allele (0, 1, or 2) in each genome to form a 2504 x 298,454 genotype matrix M. We performed PCA using the R function call prcomp(M, center=TRUE, scale=TRUE).

### Evaluation of fingerprints for population assignment

We computed a population fingerprint for each of the annotated populations studied by the 1000 Genomes Project. To identify the population closest to each individual, we compared each individual’s normalized fingerprint to the population fingerprints (using Spearman correlation, as described for comparing individual fingerprints). Each individual was considered classified as belonging to the closest population. To avoid distortions, we excluded each individual from the computation of their own population fingerprint.

## Results

### A method for encoding genotyping data

We developed an algorithm for computing ‘fingerprints’ from genotype data, including data produced by DTC genetics companies, e.g., 23andMe. Like our previously published genome fingerprinting method (Glusman et al., 2017), the new algorithm is based on locality-sensitive hashing (Indyk and Motwani, 1998). The genotype fingerprints are generated efficiently, only need to be computed once per individual, and can be efficiently compared to determine whether two genotypes represent the same individual, closely related individuals, or unrelated individuals. As with genome fingerprints, the original data cannot be reconstructed from the genotype fingerprint, enabling sharing of fingerprints for comparison when privacy concerns prevent sharing the genotype file itself.

Importantly, the genotype fingerprints can be computed starting from any of several chip array designs, with genome coordinates expressed relative to any reference version, and the resulting fingerprints are directly comparable without further conversion. We give examples using SNP lists derived from two array designs used by 23andMe: V2, based on Illumina HumanHap550 Genotyping BeadChip (~550,000 SNPs) and V3, based on Illumina OmniExpress Genotyping BeadChip (~960,000 SNPs).

The main parameter of our algorithm, *L*, determines the size of the fingerprint. Smaller fingerprints (e.g., *L*=100) are useful for fast genome comparisons to determine identity, while larger fingerprints (e.g., *L*=1000, 5000) retain more information and better support detailed analyses like population reconstruction.

### Computation on genotype fingerprints is fast

Computation of raw genotype fingerprints is very efficient, typically requiring 10-15 seconds per genome, depending on the number of genotypes included in the hybridization array design. The computation requires a single-pass read of the genotype file and depends principally on the time it takes to read the file (I/O bound). It is also trivially parallelizable: computation of one genotype fingerprint does not depend on the results of similar computations for other individuals. Computing population fingerprints for the 26 populations of the 1000 Genomes cohort took under a minute; fingerprint normalization averaged 0.13 seconds per genome; serializing the 1000 Genomes dataset into a searchable fingerprint database then took 37 seconds. Finally, all-against-all comparisons in this data set (3,133,756 comparisons) took 15 CPU seconds for *L*=1000 (4.8 microseconds per comparison) and 79 CPU seconds for *L*=5000 (25.2 microseconds per comparison). As for genome fingerprints, comparisons of genotype fingerprints are independent and trivially parallelizable.

### Fast relationship detection

We computed pairwise correlations of normalized genotype fingerprints within a family of five who made their 23andMe genotype results publicly available (Glusman et al., 2012). The resulting correlation levels are consistent with the known family relationships (Figure 2). The genotype fingerprint correlations among full siblings (Aunt and Mother, Daughter and Son) and parent-offspring pairs are higher than observed in avuncular relationships (Aunt and Daughter, Aunt and Son); unrelated pairs (Aunt and Father, Mother and Father) showed the lowest correlations. The correlations between the Son and the other family members were somewhat reduced due to comparison across different SNP lists (V2 for Son, V3 for all others).

**Figure 2.**
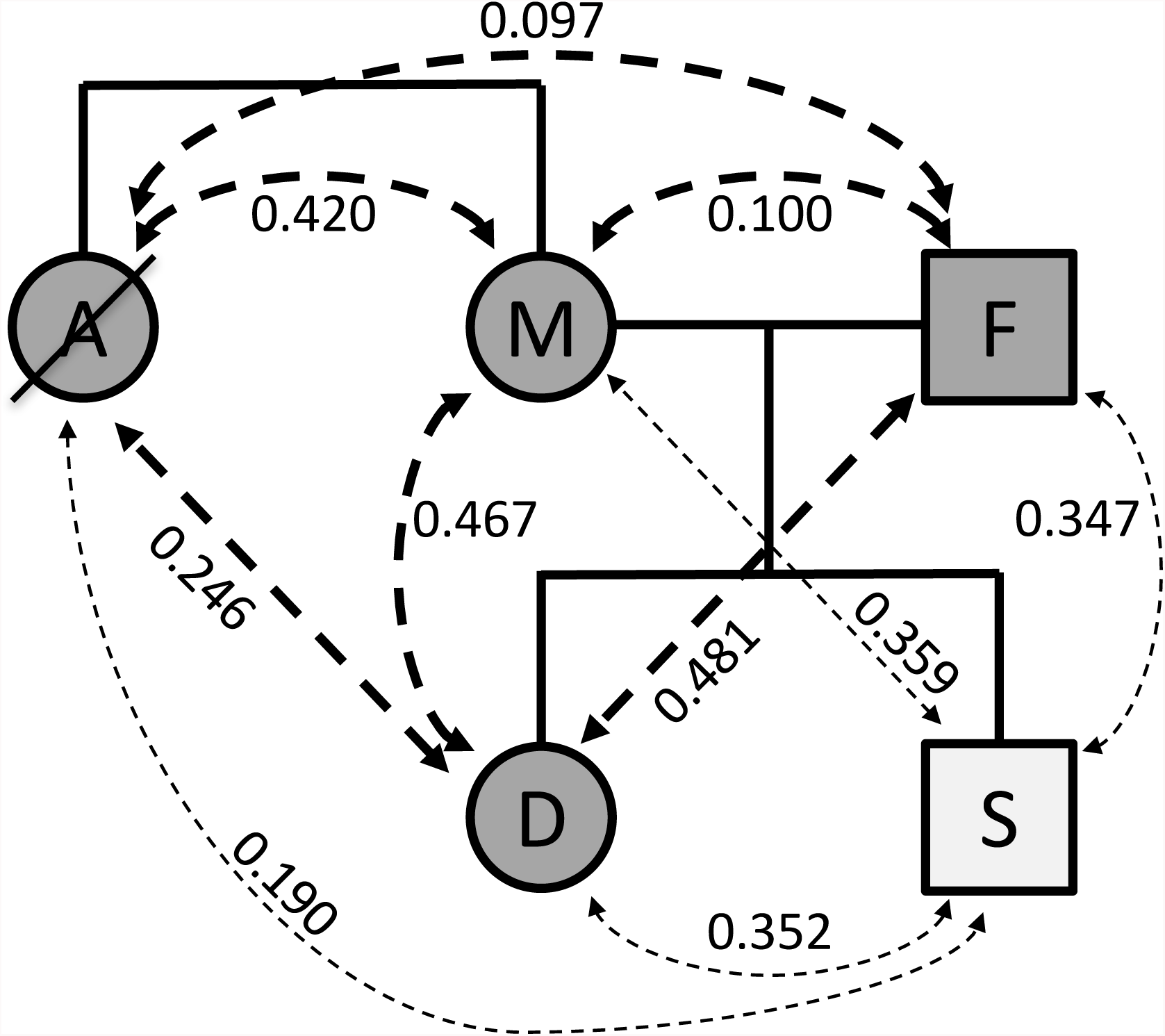
Comparison within a family of five. A: Aunt (deceased); M: Mother; F: Father; D: Daughter; S: Son. Dashed lines represent family relationships; thin lines denote comparison between individuals assayed on different versions of the genotyping platform.

### Fast analysis of population structure

We tested the utility of genotype fingerprints for population studies. We deduced genotypes for the 2504 individuals of the 1000 Genomes Project cohort using the 23andMe V3 SNP list, computed genotype fingerprints for each, and used PCA to reconstruct the known population structure (Figure 3) in a fraction of the time required to perform the same task using standard methods, and with much smaller memory requirements. The quality of the reconstruction depended on fingerprint size: fingerprints with *L*=5000 yielded excellent population structure reconstruction, comparable to the results of population reconstruction using high-resolution genome fingerprints (Glusman et al., 2017). Genotype fingerprints with smaller values of *L* progressively yielded lower-resolution results but with very significant gains in speed. As for genome fingerprints, the protocol for reconstructing populations is much simplified using genotype fingerprints: it is possible to reconstruct population structure by computing fingerprints directly from the individual genotypes and combining the fingerprints into a matrix ready for analysis with PCA, t-SNE, etc.

**Figure 3.**
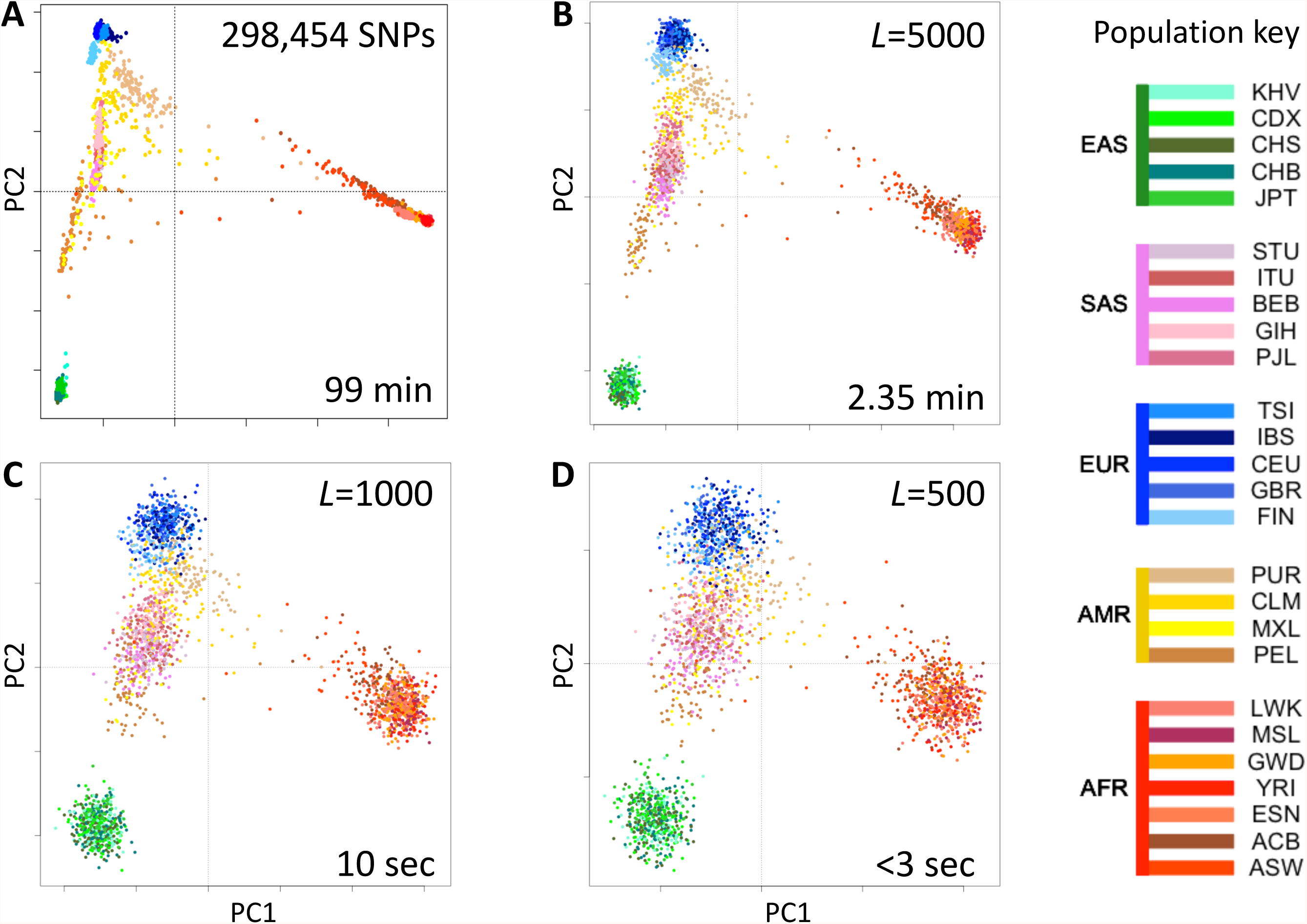
Estimates of population structure in the 1000 Genomes Project data set at different resolutions. Individuals are color coded according to their population as per the key to the right. EAS, SAS, EUR, AMR and AFR: East Asian, South Asian, European, Admixed American, and African, respectively. (A) Principal components analysis (PCA) of the 2504 individuals using ~300,000 SNPs. (B) PCA on genotype fingerprints with *L*=5000. (C) PCA on genotype fingerprints with *L*=1000. (D) PCA on genotype fingerprints with *L*=500.

### Fast population assignment

We computed “population fingerprints” in the 1000 Genomes data set by averaging genotype fingerprints (V3 set, *L*=5000) of the individuals in each population. To determine each individual’s population of origin, we then compute the correlation between the fingerprint of a query genome and the fingerprint of each population; the individual fingerprint is simply assigned to the population with which it is most strongly correlated. We tested this method by “leave one out” cross-validation. The correct population was identified as the best match for 2027 of 2504 query genomes (81% of cases). Accepting the 2nd or 3rd best population matches increased the success rate to 92.9% and 96.1%, respectively. As expected, classification was most difficult for the admixed AMR genomes; excluding these increased the success rate to 85.7%, 97.8% and 99.2% for 1st, 2nd and 3rd population matches, respectively.

### Robustness to SNP list

We evaluated whether genotype fingerprints can be compared across chip array designs. We computed genotype fingerprints for the 2504 individuals in the 1000 Genomes Project data set, to match the chip array designs used in 23andMe V2 and V3 (548,911 and 902,448 SNPs, respectively), yielding a mixed set of 5008 genotype fingerprints (two fingerprints for each of 2504 individuals). We studied this joint set using PCA (Figure 4A) and observed that the first two principal components reconstruct the known population structure (as in Figure 3); the third principal component separates between fingerprints computed on V2 and V3 versions (Figure 4B). Furthermore, the correlations between the two versions of each individual are always higher than those between related individuals (PO, FS), which are in turn higher than those between unrelated individuals (Figure 4C).

**Figure 4.**
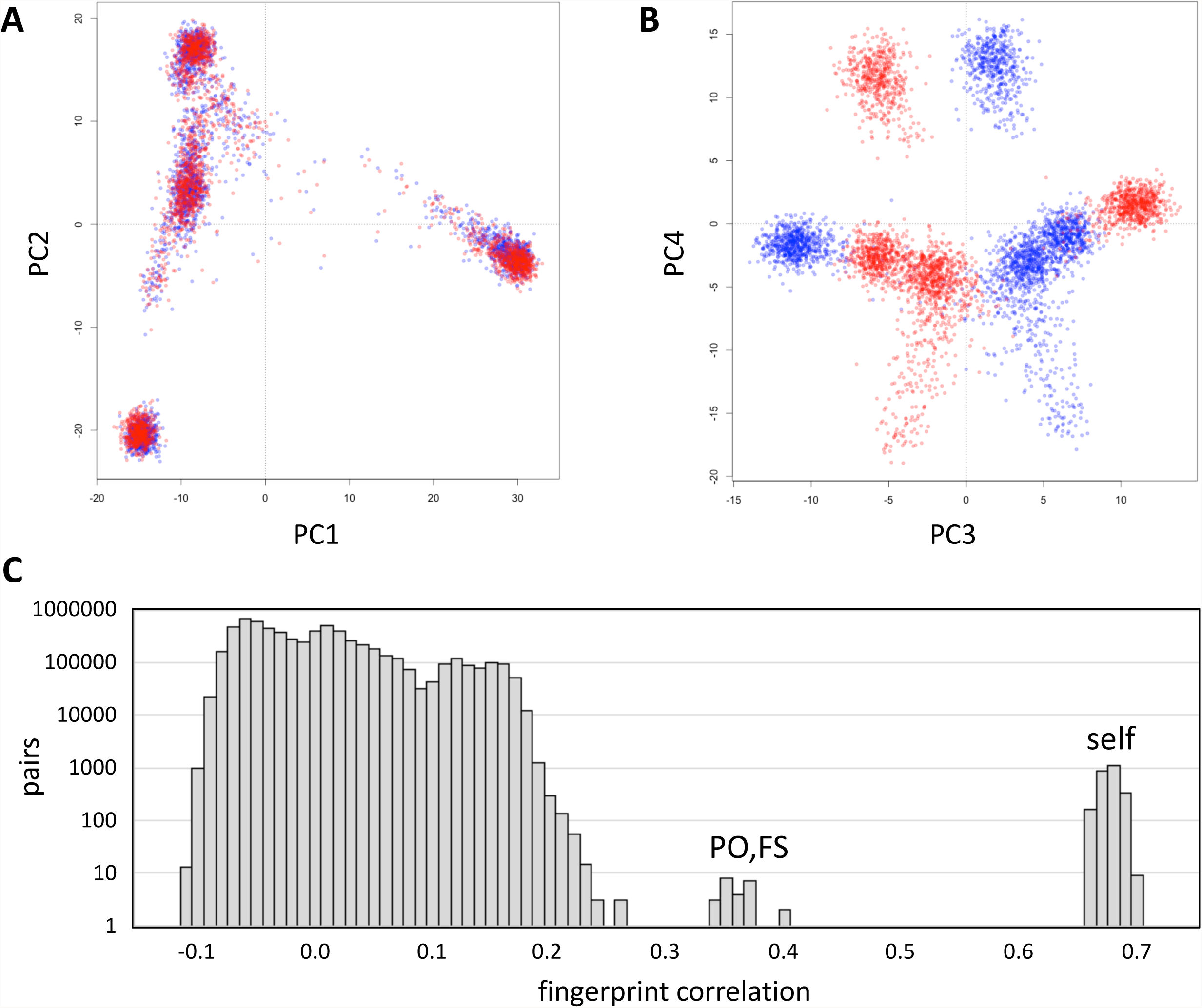
Comparison of genotype fingerprints relative to different SNP lists. We deduced normalized genotype fingerprints (L=5000) for the 1000 Genomes Project cohort using the 23andMe V2 (red) and V3 (blue) SNP lists. (A) First two principal components, showing population structure. (B) Third and fourth principal components, showing separation between the two SNP lists. (C) Distribution of cross-correlations between the two sets of genotype fingerprints (all possible pairs of V2 vs. V3). Comparisons between the two genotype fingerprints for the same individual (self) and comparisons between parent/offspring and full-sibling pairs (PO, FS) formed distinct, high-correlation subsets.

### Fast detection of close relationships

Population adjusted genotype fingerprints allow analysis of relationships within a population, increasing the resolution of close relationship analyses. We adjusted the *L*=5000 fingerprints for the 2504 individuals in the 1000 Genomes data set, matching 23andMe’s V3 design, relative to their stated population of origin, performed all pairwise comparisons, compared with kinship coefficients computed using KING (Manichaikul et al., 2010) and with previously reported relationships (Gazal et al., 2015) computed using RELPAIR (Epstein et al., 2000) (Figure 5). The similarity between unrelated individuals derived from the same population (e.g., Figure 3) is removed by adjustment to the population average. Thus, population-adjusted fingerprints for unrelated individuals show no significant correlation. As with genome fingerprints, this comparison of population-adjusted genotype fingerprints allowed the detection of related individuals in the 1000 Genomes cohort, consistent with previous reports (Gazal et al., 2015). The highly-correlated pairs correspond to a variety of degrees of relationship, from full siblings to cousins. For related pairs, fingerprint correlation levels are correlated with KING kinship coefficients. Parent/offspring and full sibling relationships, which have the same expected KING kinship coefficient (0.25) but different variance from that expected value, produced equivalent high fingerprint correlations.

**Figure 5.**
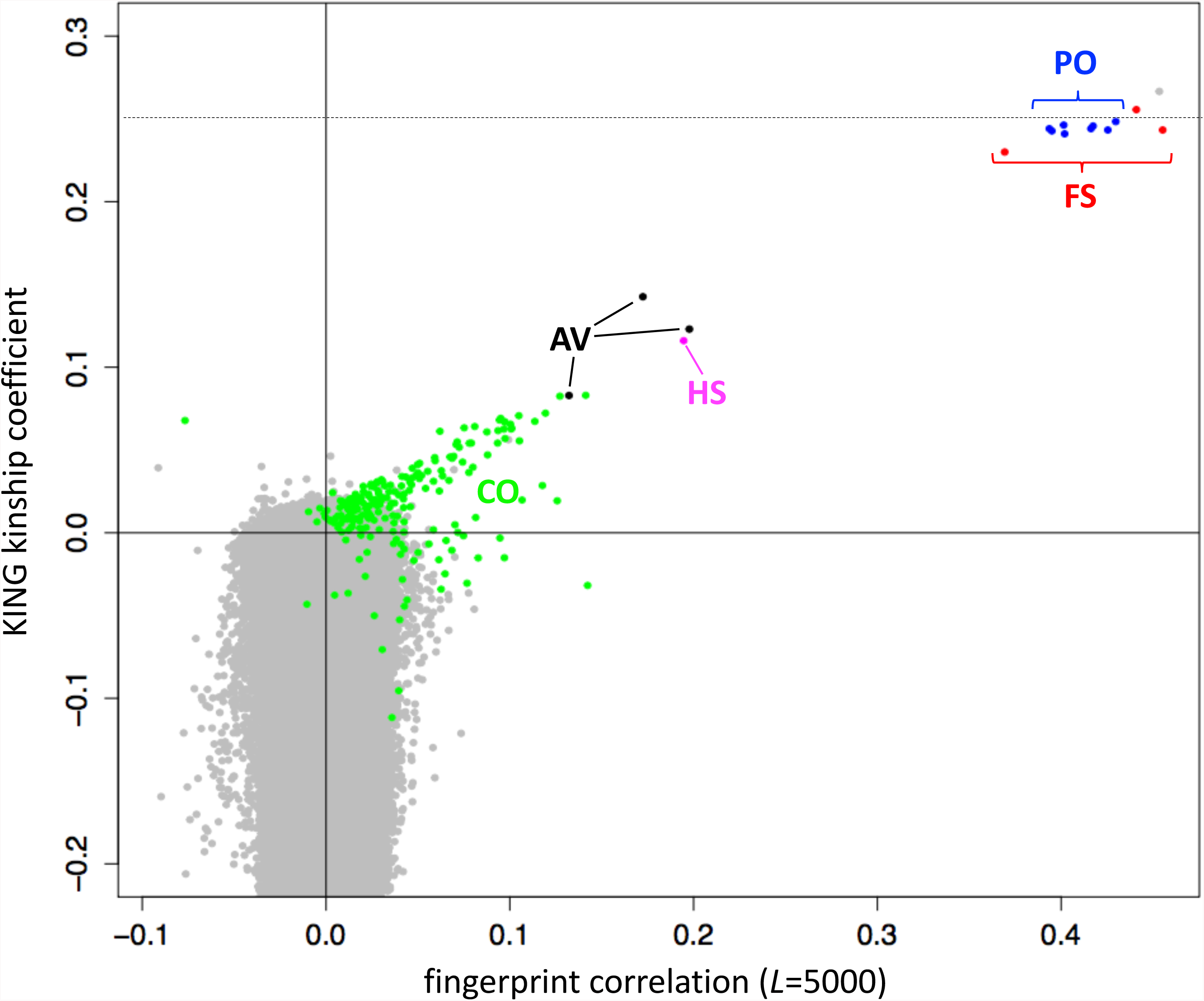
Identification of close relationships in the 1000 Genomes Project. Comparison between the correlations of population-adjusted genotype fingerprints (V3 set, *L*=5000) and the kinship coefficient as computed using KING, highlighting close relationships identified using RELPAIR. FS: full siblings (red). PO: parent/offspring (blue). HS: half siblings (magenta). AV: avuncular (black). CO: cousins (green). All other pairs in gray. One FS pair (HG03873 and HG03998, with maximal kinship, in gray) was not identified by RELPAIR.

## Discussion

We presented a method for computing ‘fingerprints’ of genome-wide SNP array genotypes as reported by DTC genetics companies, focusing on 23andMe as example. Like our previously reported fingerprints from whole-genome resequencing data, genotype fingerprints retain sufficient information to enable ultrafast comparison of genotypes, without revealing the private, detailed genotype data necessary to predict phenotypes.

We demonstrated the utility of genotype fingerprints for rapid versions of common tasks: identifying genotypes from the same individual, from closely related individuals, or from a known population, and to cluster individuals into sub-populations *de novo*. Based on comparison between two different array designs (23andMe V2 and V3), genotype fingerprints are robust to differences in number of SNPs assayed, both for establishing identity or relatedness between two samples and for reconstructing populations.

Genotype fingerprints are an adaptation of our previously published genome fingerprinting method (Glusman et al., 2017) to the more limited information present in a genotype. While genome fingerprints encode consecutive pairs of SNVs using nucleotide sequence and separation distance, genotype fingerprints encode individual SNPs using alleles, allele frequencies, and rsids. SNPs are SNVs with high population frequency; while additional SNVs are discovered as further genomes are sequenced, the vast majority of SNPs have already been identified and assigned stable identifiers (rsids). In contrast, many SNVs either lack identifiers or have been assigned preliminary identifiers that are subject to change (e.g., by merging with a different identifier representing the same variant). Stable identifiers facilitate matching variants across genome reference versions, enabling the desired robustness to a changing reference genome using the simpler encoding method presented here.

Insertions and deletions can also be represented in genotype files (e.g., using the symbols I and D, respectively) but they are much less common than SNPs and may not be present when deriving genotype files from WGS or exome data. We therefore chose to exclude them for the sake of simplicity and consistency in analysis, in similarity with the procedure we established for computing genome fingerprints from VCF files. We also chose to include only autosomal SNPs, as inclusion of variants in the sex chromosomes may lead to distorted similarity values.

There are several privacy considerations in sharing genomic information. Much attention has been paid to the risk of re-identification of de-identified samples (Ehrlich et al., 2014), even when querying genetic data sets via bandwidth-limiting interfaces like the GA4GH beacons, giving rise to strategies such as obscuring rare variants and budgeting queries (Raisaro et al., 2017). While enabling an important and powerful query - namely, “has this allele been seen before?” - these strategies for preventing re-identification preclude multiple other potential applications, thus limiting the utility of genome data sharing. Some genome data sharing scenarios exist in which anonymity is not an issue, but phenotype prediction is. For example, an individual may wish to compare their genotype (obtained via a DTC genetic testing company) to the genotypes of other individuals, without revealing to others whether they carry alleles that confer risk to develop a specific phenotype, e.g., Alzheimer’s disease - both currently known alleles, and ones whose significance may be discovered in the future. Like genome fingerprints, genotype fingerprints decouple genotype comparison from genotype interpretation, supporting the identification of closely related individuals, without exposing variant information.

The number of private individuals who already have knowledge of their genotypes, as assessed by DTC genetics companies, vastly exceeds the number of individuals with full genome data. We expect genotype fingerprints to have immediate applicability for facilitating genotype comparisons, empowering citizen science without concomitantly revealing sensitive private genetic information.

## Internet Resources

a. http://haplogroup.org/exploring-microarray-chips/
b. https://isogg.org/wiki/List_of_DNA_testing_companies
c. http://dx.doi.org/10.6084/m9.figshare.92682
d. Documentation, code, and sample datasets are available at:
  - http://db.systemsbiology.net/gestalt/genotype_fingerprints
  - https://github.com/gglusman/genotype-fingerprints

## Author contributions

GG designed the study. MR and GG performed analyses, wrote the manuscript and approved its final version.

## Acknowledgements

We wish to thank the Corpas family for releasing their individual genotypes as CC0. This work was supported by NIH grant U54 EB020406.

